# Structural motif search across the protein-universe with Folddisco

**DOI:** 10.1101/2025.07.06.663357

**Authors:** Hyunbin Kim, Rachel Seongeun Kim, Milot Mirdita, Jaewon Yoon, Martin Steinegger

## Abstract

Detecting similar protein structural motifs, functionally crucial short 3D patterns, in large structure collections is computationally prohibitive. Therefore, we developed Folddisco, which overcomes this through an index of position-independent geometric features, including side-chain orientation, combined with a rarity-based scoring system. Folddisco’s compact 1.45-terabyte index of 53 million AFDB50 structures enables querying this database within seconds. It is 20-fold faster in querying and 4-fold more storage-efficient than existing methods. Folddisco is more accurate and storage-efficient than state-of-the-art methods, while being an order of magnitude faster. Folddisco is free software available at folddisco.foldseek.com and a webserver at search.foldseek.com/folddisco.

---

Structural motifs are short, recurring arrangements of tertiary structural elements that form recognizable patterns in proteins and are often associated with stability, binding interactions, or active sites (1, 2). Evolutionary constraints due to motif functionality often result in their conservation at sub-Angstrom resolution. Thus, identifying these motifs can provide functional insights, even for proteins with unknown function (3). Notable examples for the links between structural motifs and function include the Cys2-His2 zinc finger motif in transcription factors and the CWxP, NPxxY and DRY motifs in G-protein-coupled receptors (GPCRs), which have been shown to bind zinc ions, thereby stabilizing DNA binding structure (4) and to be involved in receptor activation (5), respectively. Although structural motifs can directly provide functional insights, most functional annotation methods rely predominantly on sequence information, despite its indirect relationship to function (6). This is largely due to the high throughput of sequencing and alignment techniques (7, 8), in contrast to the relative scarcity of structural data and the limited capabilities of structure-comparison methods until recently.

However, recent revolutionary advances by AlphaFold2 (9) and other deep learning-based structure prediction methods now offer hundreds of millions (10, 11) of protein structures. These advances have motivated the development of rapid and scalable structural aligners, such as Foldseek (12), which exploits this potential by enabling direct structure-based functional annotation (13). Despite its strengths, Foldseek is not built for motif detection, as it assumes that residues match in linear order, in contrast to the non-linear path of far-apart matching pieces, common to structural motifs.

The RCSB motif search method (14) tackles the non-linearity problem by breaking each structure into proximal residue pairs. It extracts for each pair, the residues’ amino acid (AA) identities as well as geometric features: the distance between their Cα atoms, the distance between their Cβ atoms, and the intersecting angle between the Cα-Cβ vectors. Each such 5-feature set is saved in an inverted index, which maps it to the PDB entry and positions where it occurs.

Since the number of proximal pairs scales roughly with each structure’s residue count, indexing requires ∼ 75x more feature extraction and storage operations than the number of residues. As a result, the RCSB method took 3.5 days and 55GB to index 160,467 structures from the Protein Data Bank (PDB) (15). pyScoMotif (16) is a faster Python-based motif finder utilizing the same pair representation, except that it uses side-chain centroids instead of Cβ atoms. It reduced the indexing time to 20.5 hours for 195,000 structures, requiring 73GB storage space, making these the limiting factors.

Another limitation of current motif search methods is their lack of flexibility in handling various query motif types and lengths. For instance, RCSB’s service supports query motifs of up to 10 residues, restricting the method only to short motifs. Alignment-based fragment search methods, like MASTER (17), can handle longer, discontinuous queries, but struggle with short motifs like catalytic triads or zinc fingers.

Here, we present Folddisco (Fig. 1a-c), the first motif search algorithm that supports querying both short motif queries (Fig. 1d) and long, discontinuous segments (Fig. 1e) in seconds against 53 million structures. It operates efficiently on a massive scale, indexing these in under 25 hours (<1.5 TB), 11x faster to build and 4x smaller than the state-of-the-art.

**Fig. 1.**
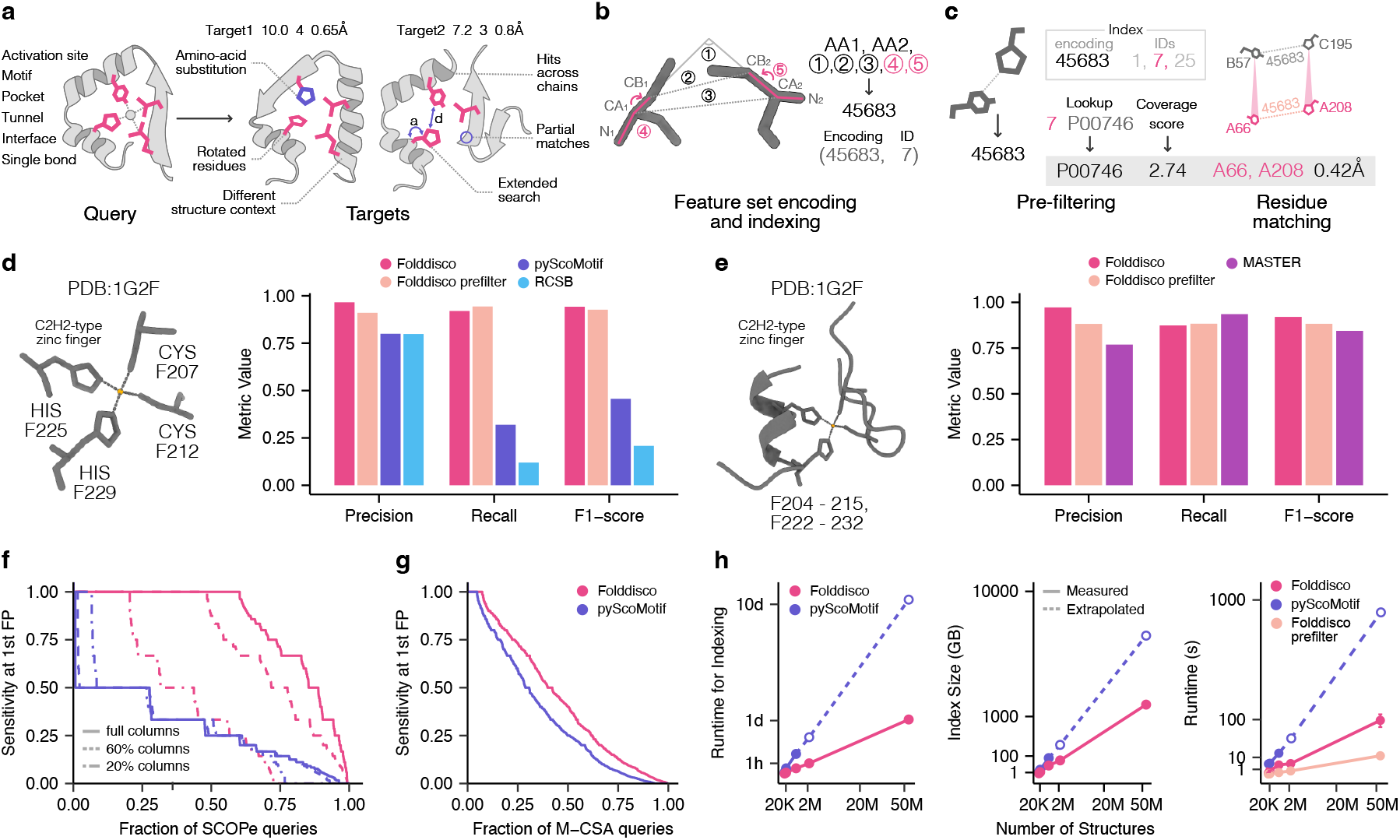
Folddisco’s workflow and benchmark. **a**, Overview: Folddisco is a fast tool for sensitive motif detection in millions of protein structures. Given motif-defining query residues (a, left), it can detect full (a, left target) or partial matches (a, right target) by comparing feature sets (b) computed from their pairs of proximal residues (<20Å). To increase sensitivity, Folddisco can compute additional query feature sets accounting for amino-acid substitutions and increased distances or angles (Methods “Extended search”). Each set is encoded (b) and rapidly searched (c) against a precomputed index of encoded feature sets from database structures. **b**, Folddisco extracts a set of 5 RCSB features (black) and 2 new features (pink) from pairs of proximate residues and bit-encodes it. Each target structure is associated with an ID and stored in an index that maps encoded feature sets to the IDs in which they are found. **c**, Querying: query feature extraction and encoding (left) is followed by retrieval of structure IDs that share its feature sets (“pre-filter”). Pre-filtered structures can be further processed to match their residues (pink) to the query (gray). **d**,**e**, Folddisco is the most accurate method in querying the human fraction of the AFDB-proteome for zinc fingers, both when using a short motif query suitable for pyScoMotif and RCSB (**d**, left; residue labels, e.g. F207, denote chain and residue number) and when using the motif-containing segments suitable for MASTER (**e**, left). **f** SCOPe-constructed benchmarks, where the goal is to match SCOPe40 sequences of the same family as the query before matching a different fold, using all conserved columns (“full”) or a random subsample of them (60%, 20%); (**g**) M-CSA hand-curated catalytic sites benchmark, counting detected PDB structures annotated as homologs as true positives. **h**, Scalability comparison on various sized databases. Indexing speed (left): Folddisco is 11x faster than pyScoMotif; Index size (middle): Folddisco’s index is 4x smaller than pyScoMotif’s; Querying speed (right) was measured for 886 (755) catalytic-site motif queries for Folddisco (pyScoMotif). The median runtime is depicted as a full point, with whiskers indicating the 1st and 3rd quantiles. Extrapolated values appear as empty points.

Folddisco examines proximal residue pairs in each input structure, extracts a set of features from each pair, encodes the set numerically, and stores it in an index (Methods, Fig. 1b). Extending the RCSB’s feature set, Folddisco introduces two additional features: the torsion angles between the N–C*α* and C*β* atoms, used by trRosetta for structure prediction (18). Folddisco generates a compact, position-less index that maps features only to structure IDs and takes advantage of the sparsity of the feature space (Methods, Supp. Fig. 1). Furthermore, the two torsion angles capture side-chain orientation and improve search accuracy (Methods, Supp. Fig. 2).

Folddisco’s full querying pipeline consists of four steps: feature extraction, pre-filtering, residue matching, and superposition (Fig. 1c, Supp. Fig. 3). First, the query motif’s feature sets are extracted and encoded as integers. Then, Folddisco pre-filters structures that share at least one feature set with the query motif, including variants within defined distance, angle, and amino acid tolerances, by looking up the query’s encodings in the index. Folddisco prioritizes the most relevant candidate structures by introducing a coverage score to rank them by their specificity to the motif. This score (Methods) uses Inverse Document Frequency (19, IDF) weights computed over the entire index, rewarding rare feature sets and penalizing common ones (e.g., helices). A length penalty further reduces random hits in large proteins, ensuring consistency for queries ranging from a single residue pair to an entire structure.

Next, Folddisco identifies the motif-forming residues in each candidate that passed the pre-filter. To do so, it constructs a graph where each candidate residue is a node, and directed edges are drawn between node pairs that match the query motif’s residue pairs either by having the same feature set (as detected in the pre-filter) or a similar one (extended search, see Methods). Folddisco detects connected components in this graph, each of which is a proposed match to the motif that is superposed to it, and the match’s root mean square deviation (RMSD) is computed.

We compared the accuracy of Folddisco with that of RCSB and pyScoMotif to detect the zinc finger motif (partial or full) and the serine peptidase motif in 23,391 AlphaFold2-predicted structures of the human proteome (Methods). Each detection of the zinc finger motif was counted as true positive (TP) if it matched the PROSITE (20) rule PRU00042, and as false positive (FP) otherwise. For serine peptidase, TPs belonged to MEROPS’ (21) family S1 and other detections were considered as FPs. Precision, recall, and F1 scores were calculated using these counts and the total number of positives (P) for the zinc finger (P=761) and serine peptidase (P=124) motifs in the human proteome. All three tools performed comparably on serine peptidase and the partial (three residues) zinc finger motif (Extended Data Fig. 1). However, Folddisco outperformed both methods when querying the full, four-residue zinc finger motif, where RCSB and pyScoMotif had low recall (Fig. 1d). Folddisco was also more accurate than MASTER (Fig. 1e) when the query was provided as segments containing the zinc finger motif (Methods), while running queries 7 times faster (full pipeline), 1,730 times faster (prefilter-only), and using a 10-fold smaller index. On this segmented input, pyScoMotif failed to return any results (Supp. Table 1). Thus, Folddisco is the only method that can search both discrete motifs and discontinuous segments. In all benchmarks, Folddisco’s prefilter alone achieved competitive accuracy, demonstrating the effectiveness of its extended feature set and coverage score ranking. We studied the runtime and scalability of Folddisco in a separate benchmark but even in this one with a single database, Folddisco’s query time was >10 times faster than any other method (Extended Data Fig. 1). To evaluate Folddisco’s generalizability beyond zinc fingers and serine peptidases, we developed a benchmark based on SCOPe40 (22), the SCOPe database clustered at 40 % sequence identity. Instead of relying on full SCOPe-domain alignments, we mimicked motifs by selecting conserved and scattered residues from family-level multiple structure alignments generated by FoldMason (23).

In each alignment we identified columns with full occupancy (no gaps) and a “dominant” residue (occurring in >66% of the members). Restricted to alignments with at least 10 such columns, we constructed three benchmarks by using the dominant residues from all identified columns as a query (termed “full”); by randomly sampling 60% of the columns and using their dominant residues; and by sampling and using 20%. This resulted in 5,753 queries (see Methods), each of which was searched against the SCOPe40 database with 15,177 structures. A match was counted as TP if it belonged to the same family as the query, FP if it belonged to a different fold, and ignored otherwise. Sensitivity was measured as the fraction of correctly identified family members (TP/P) before the first FP, where the ranking of the matches was by the coverage score for Folddisco’s pre-filter or by RMSD for pyScoMotif and Folddisco’s full pipeline. Folddisco was consistently more sensitive than pyScoMotif with Area Under the Curve (AUC) values of 0.837, 0.732 and 0.504 compared to 0.285, 0.290, and 0.300 for the three benchmarks, respectively (Fig. 1f). Of note, while pyScoMotif’s performance peaked when queries had fewest residues (on 20%) and was nearly the same otherwise (60% and full), Folddisco improved with every gain of information. In contrast to pyScoMotif, which only reports full-length matches ranked by RMSD, Folddisco also identifies partial hits. Its default IDF-based coverage score performs robustly across various motif lengths and partial matches. However, for short motifs, grouping hits by query residue coverage and ranking them by RMSD further improves sensitivity (Supp. Fig. 4). For longer motifs, RMSD ranking is less effective, as it cannot meaningfully compare partial matches across different coverage groups. Since Folddisco’s coverage score is computed entirely during the pre-filter step, it achieves identical sensitivity to the full pipeline while being approximately 10-fold faster than the RMSD-based ranking.

Complementing our synthetic structural family analysis, we assessed the detection of conserved functional sites by bench-marking Folddisco against pyScoMotif using the Mechanism and Catalytic Site Atlas (M-CSA) dataset, which provides hand-curated reference catalytic sites and annotated homologs (24). Folddisco (default) and pyScoMotif achieved AUCs of 0.432 and 0.344, respectively; a 25.6% improvement by Folddisco (Fig. 1g). With a “Sensitive” parameter setting (Supp. Fig. 5, see Methods), Folddisco reached an AUC of 0.463.

Next, we studied Folddisco’s speed and scalability in comparison to pyScoMotif, a locally runnable re-implementation of the RCSB motif search that uses the same pairwise feature set. To that end, we used Folddisco to index five databases, holding between 4K and 53M structures (Methods) and pyScoMotif to index the three smallest of them. Index construction by Folddisco was faster than by pyScoMotif, taking 18 minutes for 540K structures using 64 cores, compared to 3.46 hours (Fig. 1h, left). Folddisco’s storage requirement of 23.2GB was less than one third of pyScoMotif’s 79GB for the 540K database (Fig. 1h, middle). By extrapolating the requirements of pyScoMotif for larger databases, we find that Folddisco’s index construction time and storage requirement improve even more as the input database increases. This means, for example, that indexing the 53M structures of the AFDB50 required 1.45TB by Folddisco, compared to 4 times more for pyScoMotif (5.7TB extrapolated). Using the indices created by each tool, we measured the query runtime of Folddisco and pyScoMotif on 886 and 755 catalytic-site motifs, respectively (Methods; Supp. Table 2 and 3). Folddisco’s full pipeline was 20 and 18 times faster than pyScoMotif for indices of 20K and 500K, respectively, while Folddisco’s pre-filter only was 130 and 86 times faster on the same indices (all speedups represent median values). Folddisco’s pre-filter was nearly instantaneous for smaller databases and taking only ∼ 12 seconds for AFDB50 (Fig. 1h, right).

Having established Folddisco’s motif search capability, we examined three use cases: functional annotation of divergent sequences, search for protein state-defining motifs and interface detection. Folddisco identified a zinc finger motif in metagenomic-(from ESM30) and uncharacterized oyster (from AFDB50) proteins, which lack sequence-level annotations such as InterPro (8) domains (Fig. 2a, left and middle). It also recognized a partial motif in *E. coli*, pinpointing known metal-coordinating sites (25) of peptide deformylase (Fig. 2a, right). These examples demonstrate the advantage of Folddisco over Foldseek for detecting motifs, rather than longer structural elements, as Foldseek scored the two uncharacterized proteins with high E-values of >20 (commonly above the cutoff for discarding) and could not at all align the query to the *E. coli* protein. These discoveries and similar ones (Supp. Fig. 6) underscore Folddisco’s capacity to detect structurally conserved yet sequence-divergent features.

**Fig. 2.**
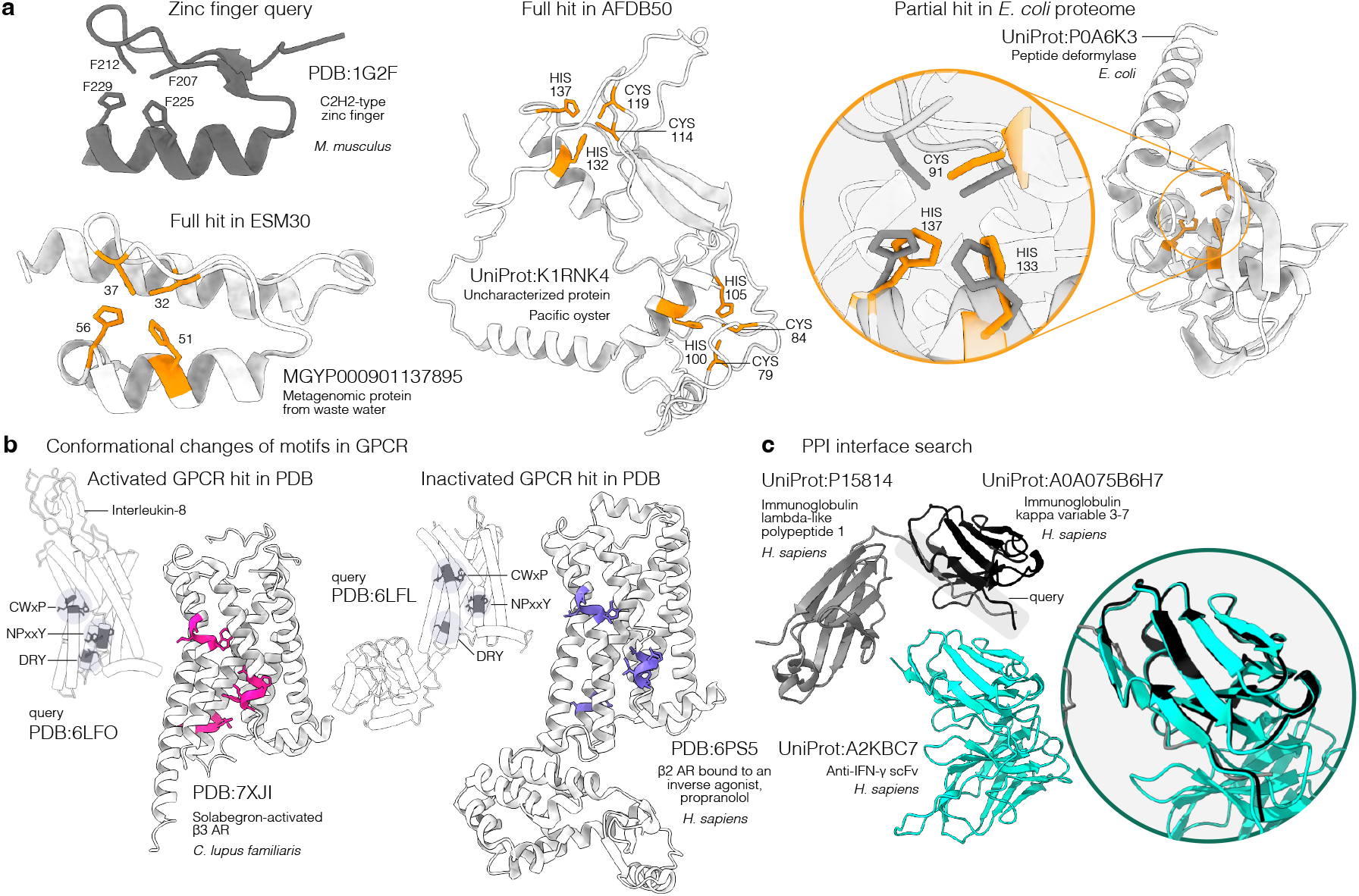
Applications of Folddisco. **a**, Zinc finger motif detection. A query for a C2H2 zinc finger (top-left) identifies full hits in previously unannotated proteins (bottom-left, middle) and a partial hit corresponding to the known metal-coordinating site in an E. coli enzyme (right). **b**, Conformational state identification. Queries using motifs from activated (left, magenta) or inactivated (right, purple) G-protein-coupled receptors (GPCRs) successfully retrieve structures in the corresponding functional states. **c**, Protein interface search. A query using an immunoglobulin domain interface (left) retrieves a single-chain variable fragment that exhibits a similar binding geometry (right).

Fig. 2b demonstrates that Folddisco searches using GPCR activation motifs (CWxP, NPxxY, and DRY from CXCR2) effectively distinguish active from inactive β-adrenergic receptor structures in the PDB. Since AlphaFold2 has been shown to sample different conformational states (26), we compared the prevalence of active versus inactive states between experimental and predicted databases. When run on the PDB, 54% of Folddisco’s matches were in the active state and a similar fraction (53%) was found in the AFDB50. This suggests that the conformational landscape sampled by AlphaFold2 closely mirrors the prevalence of functional states found in the PDB.

For the last use case, Folddisco queried a cross chain protein–protein interface motif pattern (27) derived from immunoglobulin *λ*-like and *κ* variable domains (Fig. 2c) in AFDB50, retrieving a single-chain variable fragment exhibiting the same interaction geometry. Moving beyond single motifs, we tested if Folddisco could simultaneously query active and allosteric sites; of 5,893 pairs, it retrieved both sites in 82.9% and partial matches in 16.5% (Supp. Fig. 7). Demonstrating more capabilities, Folddisco successfully identified disulfide bonds (Supp. Fig. 8) and short linear motifs (Supp. Fig. 9).

To facilitate access to Folddisco motif searching, we developed a webserver (search.foldseek.com/folddisco). Queries can be provided as a standalone motif or as a full protein structure with specified motif residues. The webserver provides prebuilt indices of major databases: AFDB50, PDB, AFDB-proteome (10), ESM30, and BFVD (28). For each query, it returns up to 1,000 top-ranked matches per database, along with Folddisco scores and interactive structure visualizations. Searching for the full zinc finger motif (Fig. 1d) in all databases at once completes in approximately 100 seconds on a single core.

Folddisco’s support for short, long, and partial motifs entails trade-offs. Its connected-component residue matching may yield spurious assignments in long queries and its fixed binning width can miss borderline true positives. This connectivity constraint precludes the detection of motifs beyond 20Å, potentially missing distant functional sites like remote allosteric pockets (Supp. Fig. 7). Furthermore, the current IDF-based coverage score is suboptimal for ranking short motifs (see Supp. Fig. 4). We plan to address these limitations by implementing motif-specific E-value statistics (29) and exploring variable-binning schemes. Ultimately, we will extend support to nucleic-acid and protein-ligand motifs to analyze the full spectrum of biomolecules predicted by modern methods (30).

In conclusion, Folddisco’s compact index enhances scalability by reducing storage and indexing time, enabling fast querying of large databases like AFDB50 and ESM30. Its features for side-chain orientations and rarity-based scoring allow accurate detection of short, long and partial motifs. By uncovering motifs linked to catalysis, complex formation, and conformational regulation, Folddisco facilitates mechanistic insights across diverse taxonomic and functional landscapes.

## Supporting information

Source Data for Extended Figure 1

Source Data for Figure 1

## Methods

### General workflow

#### Indexing

Folddisco is designed to efficiently query a motif in input databases of many millions of protein structures. Therefore, we generate an index by assigning each database protein structure a numerical ID and examining its pairs of proximal residues (default radius: 20Å). From each proximal pair Folddisco extracts two sets of 7 features (see “Pairwise features”) and jointly encodes each of them as a 32-bit unsigned integer (see “Encoding feature sets as integers”). Each unsigned integer serves as a key to retrieve the input IDs of structures in which its encoded feature set was found (see “Foldisco index”). Folddisco’s index does not include the structure positions of the proximate residues, but they can be optionally reconstructed (see “Residue matching”). This strategy in combination with delta compression of the IDs results in a more compact index compared to the indices of conventional position-storing motif search methods.

#### Querying

Pairs of proximal residues are identified in the query motif in the same way input structures are processed during indexing. For each pair at positions i and j, two sets of features (i,j) and (j,i), are extracted and encoded because of feature asymmetries (see “Pairwise features”). To increase sensitivity, Folddisco can search for more encodings through amino-acid substitutions and adjustable distance/angle deviations (“Extended search”). In the pre-filter step, the 32-bit integers computed for the query motif are used as keys to retrieve IDs of indexed structures that share at least one feature set with the query. Folddisco then ranks the IDs by their coverage of the motif, i.e., by the number of feature sets they share with it and the sets’ rarity (see “Pre-filtering”). After pre-filtering, an optional step can match the residues of the query to those of pre-filtered structures. As Folddisco doesn’t index positional information, this step is conducted to compute a residue mapping by finding connected graph components (see “Residue matching”).

### Folddisco’s feature set

#### Pairwise features

Folddisco extracts two feature sets from each pair of proximal residues in each input structure as well as the query. This set includes five features used by RCSB: the amino acid type of the residues (AA1 and AA2), the distance between their Cα atoms, the distance between their Cβ atoms, and the intersecting angle between the Cα-Cβ vectors. Since RCSB’s features do not capture the side-chain orientations of the residues, we included two additional features in Folddisco’s set: the two dihedral angles in the atoms of N1-Cα1-Cβ1-Cβ2 and N2-Cα2-Cβ2-Cβ1, which were proposed by trRosetta for structure prediction (18). Since these two features are not symmetric and may have different values, depending on which residue is considered as the first, Folddisco considers two feature sets, one in the direction of AA1-AA2 and another in the direction of AA2-AA1.

#### Encoding feature sets as integers

Folddisco encodes the seven features in a set in bits and concatenates them bitwise as follows. AA1 and AA2 are treated numerically (0,1,…,19), requiring 5 bits each. The distance features are discretized into bins from 0 to 20Å (default number: 16), requiring 4 bits each. To keep the cyclic nature of the angle features, we discretize their cosine and sine values from -1 to 1 (default: 4 bins for cosine, 4 bins for sine), requiring 4 bits per angle. In total, this encoding requires 30 bits (Supp. Fig. 3a), which can be represented as a 32-bit unsigned integer (first two bits are always ‘0’).

### Foldisco index

Folddisco’s index consists of 4 files; values (no suffix), offset (.`offset`), lookup (.`lookup`), and metadata (.`type`).

The.`offset` file acts as the primary index, enabling rapid retrieval of a list of protein structure identifiers for any given feature encoding. It uses two parallel arrays: (1) a list of all observed feature encodings (keys) stored as ascending sorted 32-bit integers, and (2) a corresponding array of 64-bit byte offsets pointing into the values file. Storing only the observed encodings saves spaces compared to storing the full encodings’ space (Supp. Fig. 1) and keeping them sorted allows retrieving an entry for any key using binary search with *O*(log *N*) time complexity. Additionally, index construction can be fully multi-threaded (Supp. Fig. 10).

The value file stores the protein-structure identifiers associated with each key. To optimize storage, these identifiers are stored as delta-compressed, variable-length integers. The. `lookup` file maps these numerical IDs back to their original alphanumeric identifiers (e.g., mapping 0 to P00568), while the metadata retains query-specific details, such as input paths and binning information. Because the compressed IDs are written in parallel during index construction, we recommend using SSDs (solid state drives) to avoid input/output bottlenecks.

#### Encoding distribution analysis

The Folddisco feature is encoded with 30 bits, giving a theoretic potential of 2^30^ different encodings. A naive index implementation would reserve space for each of them *a priori*. To design a more compact index, we checked whether all encodings are seen in various databases: human proteome, AFDB Swiss-Prot, AFDB Fold-seek cluster representatives, and AFDB50. We found that most encodings (>93% in all databases) are not observed even once (Supp. Fig. 1, left-top), indicating a large compression potential, which we took advantage of in our implementation. To further investigate the difference between the theoretic and observed encodings, we computed the prevalence of both observed encodings (Supp. Fig. 1, left-bottom) as well as of co-occurring amino-acid pairs (Supp. Fig. 1, right) within the most common encodings (top 100). For all databases, we found that the most co-occurring amino-acids pairs were ALA-LEU, common in alpha helices. This suggests that helix/helix interactions dominate the index.

### Protein structure input formats

Folddisco accepts input protein structures either as files in PDB or mmCIF format (optionally gzip-compressed), or as Foldcomp-compressed (31) files. When reading Foldcomp-compressed protein structures, Folddisco can iterate through them at a comparable speed compared to reading uncompressed protein structures in PDB/mmCIF format.

### Pre-filtering

When querying a motif, Folddisco first applies a computationally inexpensive pre-filter through our index to identify candidates, eliminating most non-matching structures before any structures are read from disk (Supp. Fig. 3b). For the query motif, pairs of proximal residues are identified in the query motif (see “Extended search”), and two sets of features are extracted and encoded from each of them in the same way the input structures are processed during indexing, resulting in a set of 32-bit unsigned integer keys. Folddisco uses these keys to list candidate structures that share at least one feature set with the query motif as follows.

#### Encodings’ rarity

Folddisco computes Inverse Document Frequency (IDF) weights for each 32-bit encoding to represent its rarity among the input protein structures.

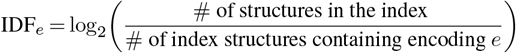

These weights are used to rank the candidate structure list by computing coverage scores.

#### Coverage scores

Each candidate is scored by the sum of the IDF weights of the encodings it shares with the query:

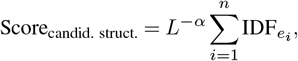

where *n* is the number of shared encodings, *L* is the length of the candidate structure in residues, and *α* is a length-penalty exponent (default 0.5) to avoid length-dependent random matches ranking high (Supp. Fig. 12).

#### Motif completeness score based on query residues

In addition to the coverage score, Folddisco computes a motif completeness score for each candidate. Notably, if a given structure candidate shares two feature sets (encodings) with the query, they can involve either 3 or 4 distinct residues in the query: the two encodings from two pairs that share a residue (x-y and x-z) or two encodings from two distinct pairs (x-y and z-t). Since Folddisco has access to the query’s residues also during pre-filtering, it computes, for each candidate, the number of distinct query residues participating in at least one feature shared between the candidate and the query. This number is reported separately from the coverage score and can be used for optional post-filtering.

#### Extended search

Based on the distance and angle features of each query proximal pair, Folddisco will produce by default up to 10 alternative feature sets. It does so by adding (or subtracting) small deviations (default = 0.5 Å for distances and 5° for angles) to each distance/angle feature, keeping the other features at their exact values. Alternative feature sets that result in invalid numbers (e.g., negative distance) are ignored and the rest are encoded as integers, as described in the previous section. Users who wish to limit the search can set the number of allowed deviation values to 0. During benchmarking, we applied default deviations to allow minor conformational differences. Folddisco also allows extended amino acid matching by providing alternative uppercase one-letter amino acid codes (e.g., ‘A’ for Alanine and ‘X’ for any amino acid) after the position of the query residue to extend. Users can also use custom lowercase one-letter codes to represent amino acid groups by properties: ‘p’ (positive), ‘n’ (negative), ‘h’ (hydrophilic), ‘b’ (hydrophobic), and ‘a’ (aromatic). Setting an amino acid alternative increases the number of computed query encodings, extending the search similarly to the distance and angle deviations.

### Residue matching and superposition

#### Graph construction

Despite not holding position information in its index, Folddisco can match the residues of an input candidate to the query after the pre-filter step by establishing a graph where the nodes represent the candidate’s residues (Supp. Fig. 3c). At this stage Folddisco reads the full information of each candidate, including residue positions from disk. Folddisco examines all candidate residue pairs that match at least one query residue pair in terms of amino acid identities (e.g., a candidate Cys-His pair will be examined if the query has Cys-His as one of its pairs). If a residue pair shares a feature set with some query residue pair, Folddisco links the residue pair’s nodes with an edge in the direction indicated by the shared feature. In this stage, more than two feature sets are considered for each query residue pair by setting the distance and angle deviations as described in the section “Extended search”. This means that the residues of all pairs discovered by the pre-filter will be linked with at least one directed edge, but potentially additional off-by-a-little pairs will be linked as well. In addition, residue pairs with the same amino acid identities and a similar Cα–Cα distance to some query residue pair (<1.0Å difference by default) will be linked with an edge.

#### Connected components detection and scoring

Folddisco lists connected components in the graph built for the candidate, as these cohesive subgraphs indicate groups (and not just pairs) of residues that together are likely to form the motif. Folddisco identifies two connectivity types: strongly connected components (SCCs) with Tarjan’s algorithm (32) and weakly connected components (WCCs) by first ignoring edge directions and then performing a DFS-based search (33). WCCs aim to capture more complete matches to the query, whereas SCCs enforce stricter residue-pair interactions. Folddisco merges the two sets of SCCs and WCCs. After residue matching, Folddisco superposes the query motif on the matched residues using the Kabsch algorithm (34). RMSD is calculated using the coordinates of the Cα and Cβ atoms of the query motif and the matched residues. To provide length-independent assessments of structural similarity, Folddisco additionally computes the Template Modeling score (TM-score) (35) and Global Distance Test scores (GDT-TS, GDT-HA) (36) between the motif matching residues of the query and target structures. To monitor outliers within matched residues, the Chamfer (37) and Hausdorff distances (38) are calculated. Complementing these geometric metrics, the IDF score is recalculated by summing up the values of the matching residues without applying length penalty.

### Benchmarks

#### Version of databases and software

We used pyScoMotif version 20231119 (commit 916b515), MASTER version 1.5, and Folddisco commit 765aeeb in all benchmarks. PDB archive was downloaded on March 7th, 2024 (209,564 entries). Catalytic site information of M-CSA and the catalog of matching homologous residues were downloaded on September 23rd, 2025. Active and allosteric sites were retrieved on Jan 22nd, 2026, from the AlloBench dataset (39), which integrated the ASD database (40) with catalytic site annotations. We used AlphaFold database version 4 (released in October 2022), ES-Matlas v0 (released in November 2022) and SCOPe v2.08. We additionally derived subsets from these resources: AFDB rep., the non-redundant representatives produced by Foldseek clustering of AFDB v4; AFDB50, the same AFDB v4 models clustered at a 50% sequence-identity threshold; and ESM30, high-confidence ESMatlas v0 models clustered at 30% sequence identity.

### Human proteome benchmark

We benchmarked Folddisco against RCSB motif search, MASTER, and pyScoMotif using the human subset of the AFDB-proteome (23,391 protein structures). Motif sets were taken from Bittrich et al. (14), including the catalytic triad of serine protease in trypsin (PDB: 4CHA, residues B57, B102, C195) and the C2H2 zinc finger motif from early growth response protein 1 (PDB: 1G2F). For the zinc finger, we tested both the three-residue partial motif (residues F207, F225, F229) and the complete four-residue motif (residues F207, F212, F225, F229). Because MASTER requires segment queries, we included neighboring residues (F204-215, F222-232) in the query. True and false positives were assessed using PROSITE annotations from UniProt (see Main).

### SCOPe benchmark

To generate motif-like queries from SCOPe40, we identified conserved residues based on family-level multiple sequence alignments. Of 15,177 SCOPe40 entries, 11,502 belonged to families with at least two members, which we aligned using FoldMason (c0430f1) with the easy-msa workflow at default parameters. From these alignments, we retained 5,753 queries that met the conservation criteria described in the main text. Folddisco searched each query using two modes: pre-filter only and the full pipeline. Folddisco successfully completed all 5,753 queries for each sampling ratio within 30 minutes. PyScoMotif searches were terminated after 8 hours; for the sampling ratios of 20%, 60%, and 100%, it completed 4,566, 5,448, and 5,315 queries, respectively. For accuracy evaluation (Fig. 1f), we considered only the queries completed by pyScoMotif.

### M-CSA benchmark

We benchmarked active-site retrieval using homologous catalytic-residue annotations from M-CSA, which are inferred via sequence homology. We constructed a set of 713 queries from M-CSA reference catalytic sites, excluding motifs with only two residues to ensure comparability with pyScoMotif. Our target database consisted of 62,122 PDB structures annotated as containing homologous catalytic sites. We measured sensitivity up to the first false positive, defining a true positive as the retrieval of a PDB structure annotated as a homolog in M-CSA, with all remaining structures in the dataset serving as negatives.

### Scalability benchmark

To assess scalability, we measured both index-construction costs and query performance across progressively larger structure collections. We built indices for pyScoMotif and Fold-disco using the proteomes of *E. coli, H. sapiens*, and SwissProt with 4,363, 23,391, and 542,378 structures, respectively. Fold-disco additionally indexed the larger datasets AFDB clusters and AFDB50 with 2,266,735 and 53,665,850 structures, respectively. Since pyScoMotif could not feasibly index the AFDB sets within a practical timeframe and space, we extrapolated its runtime and storage requirements for these collections by fitting a linear regression of those metrics against total residue count in smaller datasets. Query scalability was evaluated using catalytic-site motifs from the Mechanism and Catalytic Site Atlas (M-CSA) (41). Folddisco queried 886 annotated catalytic sites, while pyScoMotif, which requires at least three residues, was evaluated using 755 sites. All indexing benchmarks utilized 64 CPU cores, and querying tests used 12 cores.

### Folddisco parameter optimization

#### Effect of torsion angles and angle encoding

To evaluate the contribution of side-chain torsion angles and the efficacy of Folddisco’s angle encoding, we conducted two comparisons using the human proteome dataset and zinc finger (same analysis as Fig. 1d) and serine peptidase motifs (see “Benchmarks”, Supp. Fig. 2). First, we compared the default 7-feature set against an ablation setting, in which the two dihedral angles were removed, measuring both F1-score and runtime. Second, we evaluated the angle encoding strategy by comparing Fold-disco’s default method of discretizing cosine and sine values against a direct encoding of angle radians into fixed-sized bins, comparing their F1-scores.

#### Effect of binning size

We analyzed the impact of the number of bins used for feature discretization on F1-scores and runtime using the M-CSA parameter optimization subset (see “Parameter optimization”). We varied the number of bins for both distance and angle features from 4 to 32, with a step size of 4 and measured the total runtime as a function of sensitivity at the first false positive for each number of bins (Supp. Fig. 11).

#### Effect of length penalty

We investigated the effect of the length penalty on the sensitivity of pre-filtering step using the SCOPe benchmark dataset (Supp. Fig. 12). By testing penalty values ranging from 0 to 1 with a step size of 0.1, we measured the average of sensitivity up to the first false positive for each value.

#### Parameter optimization

To investigate the impact of search parameters on retrieval quality and to determine the maximum performance capability of the Folddisco motif search tool, we performed a systematic optimization using the Optuna framework (42). We constructed a representative validation subset of 45 M-CSA active sites, sampled to balance distributions across the Enzyme Commission (EC) hierarchy, residue count, and baseline retrieval difficulty (sensitivity and F1-score). Using this subset, we tuned the search parameters to maximize retrieval performance and conducted binning size trade-off experiments.

### Double motif analysis

Using the AlloBench Jupyter Notebook, we retrieved annotations and 2,146 PDB structures. We filtered for entries with fully resolved, non-overlapping active and allosteric sites, requiring at least 3 residues per site and a cumulative length ≤ 30 residues. This yielded 355 queries, which we searched against the full 2,146-structure dataset to assess Folddisco’s double-motif detection capabilities.

A total of 5,893 pairs were generated for evaluation by linking the 355 curated queries to all corresponding PDB entries sharing the same ASD Protein ID. We categorized the outcomes based on the number of matched residues (*N*_match_) relative to the sizes of the allosteric (*N*_allo_) and active (*N*_active_) sites in both the query and target annotations. A result was classified as a “DOUBLE” if *N*_match_ equaled the total number of query or target residues, or if it satisfied a dual-coverage threshold where *N*_match_ ≥ *N*_allo_ + 0.5 × *N*_active_ and *N*_match_ ≥ *N*_active_ + 0.5 × *N*_allo_. This ensures substantial recovery of both functional sites. A result was classified as a “PARTIAL” if *N*_match_ exceeded the size of a single site (*N*_allo_ or *N*_active_) but failed to meet the double motif criteria. Finally, matches were categorized as “FAIL” if they did not meet the partial threshold or if no structural match was found.

### pyScoMotif

For each dataset (*E. coli, H. sapiens*, and Swiss-Prot) a single index was built with 64 threads, and queried with 12 threads. Default scoring and filtering parameters were applied in both steps.

~~~
pyscomotif create-index pdb_dir --index_path
index --n_cores 64
pyscomotif motif-search --results_output_path
results.tsv --n_cores 12 index query.pdb
residue1 residue2 residue3 …
~~~

### RCSB’s motif search

Since the RCSB implementation provides a framework rather than a standalone method, we conducted motif searches through the RCSB web-based interface, applying the same parameters described in the pyScoMotif manuscript.

Zinc finger motifs with full residues and partial three residues were submitted with an RMSD cutoff of 1 Å and atom pairing set to “All atoms”, while restricting results to AlphaFoldDB models of *H. sapiens* (“computational” structures). For instance, the full zinc-finger motif was searched with the following query string:

~~~
QUERY: Structure Motif = 1G2F
(F_1-7 AND F_1-12 AND F_1-25 AND F_1-29)
AND RMSD Cutoff = 1
AND Atom Pairing = “All Atoms”
AND (Structure Determination Methodology = “computational”
AND Source Database = “AlphaFoldDB”
AND Scientific Name of the Source Organism = “Homo sapiens”)
~~~

### MASTER

To assess Folddisco’s segment-query performance, we used MASTER as the baseline method. First, every target structure in the `H. sapiens` proteome was converted into PDS format with `createPDS` in `--type target` mode, producing a target list for downstream searches. The zinc-finger query segment was prepared in the same way, using `--type query`. Secondly, the structural patch search was carried out with the `master` binary in single-thread mode—MASTER offers no built-in parallelization—using an RMSD cutoff of 2.5 Å (`--rmsdCut 2.5`). All other parameters remained at their defaults.

~~~
createPDS --pdbList h_sapiens_pdb_list
--pdsList h_sapiens_pds_list --type target
createPDS --pdb zinc_finger_patch.pdb --type query master --query
zinc_finger_patch.pds
--targetList h_sapiens_pds.list --rmsdCut 2.5
~~~

### Folddisco

#### Indexing

All indices were built using 64 CPU cores. For the AFDB50 subsets and ESM30, we utilized Foldcomp databases as inputs, while the PDB index was constructed from raw PDB files.

~~~
folddisco index -p db -i db_folddisco -t 64
~~~

### Motif benchmark

Searches were performed against the prebuilt *H. sapiens* index with 12 threads (`-t` 12). For zinc finger motifs, we kept structures that have at least three matching nodes from the pre-filter (`--covered-node 3`). Pre-filter-only runs were carried out with `--skip-match` to isolate the geometric hashing stage. When full matching was enabled, we filtered out structures that do not have at least one match with four residues and an RMSD of 1.0 Å (`--max-node 4 --rmsd 1.0`). Motif hits were aggregated at the structure level by enabling the `--per-structure` option. Searches for the serine-peptidase catalytic triad and for the three-residue zinc-finger core were executed with the three-node filter as well.

~~~
folddisco query -p 1G2F.pdb
-q F207,F212,F225,F229
-i h_sapiens_folddisco -t 12
--covered-node 3 --skip-match
folddisco query -p 1G2F.pdb
-q F207,F212,F225,F229
-i h_sapiens_folddisco -t 12
--covered-node 3 --max-node 4
--rmsd 1.0 --per-structure
folddisco query -p 1G2F.pdb -q F207,F225,F229
-i h_sapiens_folddisco -t 12 --covered-node 3
--skip-match
folddisco query -p 1G2F.pdb -q F207,F225,F229
-i h_sapiens_folddisco -t 12 --covered-node 3
--max-node 3 --rmsd 1.0 --per-structure
folddisco query -p 4CHA.pdb -q B57,B102,C195
-i h_sapiens_folddisco -t 12 --covered-node 3
--skip-match
folddisco query -p 4CHA.pdb -q B57,B102,C195
-i h_sapiens_folddisco -t 12 --max-node 3
--rmsd 1.0 --per-structure
~~~

#### Segment benchmark

To compare with MASTER on longer contiguous fragments, we queried two peptide segments of 1G2F using `--top 800` for the pre-filter and subsequently permitted up to 15 nodes during refinement (`--max-node 15`).

~~~
folddisco query -p 1G2F.pdb -q F204-F215,F222-F232
-i h_sapiens_folddisco --top 800 --skip-match
folddisco query -p 1G2F.pdb -q F204-F215,F222-F232
-i h_sapiens_folddisco --top 800
--per-structure --max-node 15
~~~

#### Scalability benchmark

Enzymatic active site queries from the M-CSA set were searched with a filter 75 % residue coverage within pre-filtering step and an RMSD cutoff of 2.0 Å (`--covered-node-ratio 0.75 --rmsd 2.0`). As above, each query was also repeated in pre-filter-only mode to measure the standalone cost of the residue-matching step. M-CSA queries were given as a tab-separated file with the first column containing the query structure file path and the second column containing a comma-separated residue list.

~~~
folddisco query -q mcsa_query.tsv
-i h_sapiens_folddisco -t 12
--covered-node-ratio 0.75 --skip-match
folddisco query -q mcsa_query.tsv
-i h_sapiens_folddisco -t 12
--covered-node-ratio 0.75 --rmsd 2.0
~~~

#### SCOPe benchmark

For the SCOPe benchmark, queries were executed with a retrieval limit of 200 structures (`–top 200`). We benchmarked both the full motif-search pipeline and the prefilter-only baseline.

~~~
folddisco query -q scope_query.tsv
-i scope40_folddisco -t 128
--top 200 --per-structure
--sort-by max_node_count,min_rmsd
folddisco query -q scope_query.tsv
-i scope40_folddisco -t 128
--top 200 --skip-match --sort-by idf
~~~

#### M-CSA benchmark

We evaluated Folddisco using two configurations: a standard setting (Default) and a high-sensitivity setting (Sensitive) tuned via parameter optimization. The Default profile used a retrieval limit of 6000 (`–top 6000`), while the Sensitive profile incorporated relaxed geometric tolerances and additional scoring metrics such as GDT and Chamfer distance.

~~~
folddisco query -q mcsa_query.tsv
-i mcsa_index -t 128
--top 6000
folddisco query -q mcsa_query.tsv
-i mcsa_index -t 128
-d 0.5 -a 10.0 --ca-distance 1.0
--covered-node-ratio 0.3
--max-node-ratio 0.35
--rmsd 5.0 --tm-score 0.2
--gdt-ts 0.25 --gdt-ha 0.15
--chamfer-distance 5.5
--hausdorff-distance 12.0
--sort-by node_count,gdt_ts,rmsd,idf
~~~

### Computing resource and resource measurement

All benchmarks were conducted with a server equipped with an AMD EPYC 7702P 64-core CPU, 1 TB RAM, and the same 15.3 TB NVMe storage. Maximum RAM usage (maximum resident set size) and elapsed time of each tool were measured with the GNU `time -v` command.

### Webserver

#### Querying and motif definition

We integrated Folddisco into the MMseqs2 webserver platform (43). The Folddisco motif search is available when the webserver is launched in structure mode, in conjunction with Foldseek, Foldseek-Multimer and FoldMason. Users can search through Folddisco databases for AFDB50, AFDB-proteome, ESM30, BFVD and PDB. The latter is built directly from the original PDB files, while the others were built from Foldcomp databases (Data availability). To initiate a search, the user uploads a PDB or mmCIF file containing either a motif-containing fragment or a full protein structure. By default, all residues are marked as part of the motif, but the user can refine this selection by providing or altering a motif string specifying residues of interest. To help define the motif, the interface offers two tools: an interactive motif editing widget that lets the user adjust the string graphically, and a ligand selector that parses ligands from the uploaded structure and proposes candidate motifs composed of residues within a user-defined radius (default 3.5Å) of each ligand.

#### Match reporting and exploration

The 1,000 database structures with the highest pre-filter coverage scores are reported in the webserver by using Fold-disco’s --top 1000 parameter. These are passed to Fold-disco’s residue matching step, and the final result list is then sorted by number of matched residues and RMSD. To visualize query and target residue matching, the matched target structures are decompressed from the Foldcomp database, except for PDB entries, which are retrieved directly as PDB files. All structure visualizations are rendered using the NGL viewer library (44). To help users explore the matches, the interface displays the UniProt description and taxonomic names for each entry originating from the AlphaFold databases. An integrated TaxoView (45) allows users to visualize the overall taxonomic distribution of the matches and subset results by specific taxonomic clades.

#### Identification of conserved structural motifs

The result list can be subset to show only partial motif matches. Additionally, an optional DBSCAN-based (46) clustering step groups matches with similar inter-residue distance patterns, highlighting conserved structural groups within the results. The user can adjust the clustering sensitivity by modifying the DBSCAN parameters epsilon and MinPts.

Finally, the user can download an archive containing the matches as tab-separated-values (TSV) files for each database for downstream use. Each file contains columns specifying the target accession, number of matched nodes, coverage score, RMSD, matching residues in the query, rotation and translation vectors, Cα coordinates of the matched target residue, an internal numeric target identifier, and the matched residues in the query.

## Data availability

The benchmark data is available via Zenodo at https://doi.org/10.5281/zenodo.18443780.

## Code availability

Folddisco is GPLv3-licensed free open-source software. The source code and ready-to-use binaries, as well as precomputed databases, can be downloaded at folddisco.foldseek.com. The scripts used for the benchmarks and plotting are available at https://github.com/steineggerlab/folddisco-analysis. The webserver code is available at github.com/soedinglab/mmseqs2-app.

## Acknowledgements

We deeply thank Eli Levy Karin from ELKMO, Jaebeom Kim, and Sukhwan Park for insightful discussions and their comments on the manuscript draft. This work was supported by National Research Foundation of Korea grant 2020M3-A9G7-103933 (M.S.), National Research Foundation of Korea grant RS-2020-NR049543 (M.S.), National Research Foundation of Korea grant RS-2021-NR061659 (M.S.), National Research Foundation of Korea grant RS-2021-NR056571 (M.S.), National Research Foundation of Korea grant RS-2024-00396026 (M.S.), Novo Nordisk Foundation NNF24SA0092560 (M.S.), the Creative-Pioneering Researchers Program through Seoul National University (M.S.),and National Research Foundation of Korea grant RS-2023-00250470 (M.M.).

## Author contributions

**Hyunbin Kim:** conceptualization, methodology, software, validation, formal analysis, investigation, data curation, writing-original draft, writing-review and editing, visualization, project administration. **Rachel Seongeun Kim:** conceptualization, software, investigation, data curation, writing-original draft, writing-review and editing, visualization. **Milot Mirdita:** methodology, software, data curation, writing-original draft, writing-review and editing, supervision. **Jae-won Yoon:** software, formal analysis, visualization, writing-review and editing. **Martin Steinegger:** conceptualization, methodology, software, resources, writing-original draft, writing-review and editing, visualization, supervision, project administration, funding acquisition.

---

**Extended Data Fig. 1.**
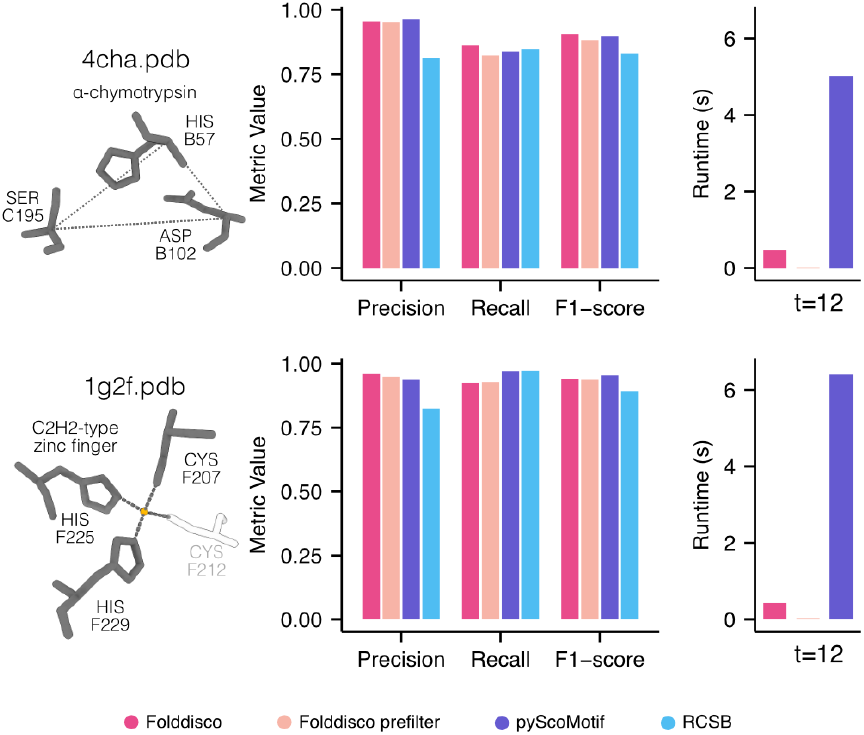
Performance comparison of Folddisco to pyScoMotif and RCSB. Precision, recall, F1-score and search time (in seconds) with 12 threads were evaluated for **top** serine peptidase motif query and **bottom** zinc-finger motif with 3 residues. Folddisco achieved the best performance for the serine peptidase query. While pyScoMotif and RCSB exhibited low sensitivity on the 4-residue zinc-finger query, they performed well with 3 residues, where Folddisco showed comparable performance (ranking second). Consistently, Folddisco was over 10 times faster than pyScoMotif across these settings.

**Supplementary Figure 1.**
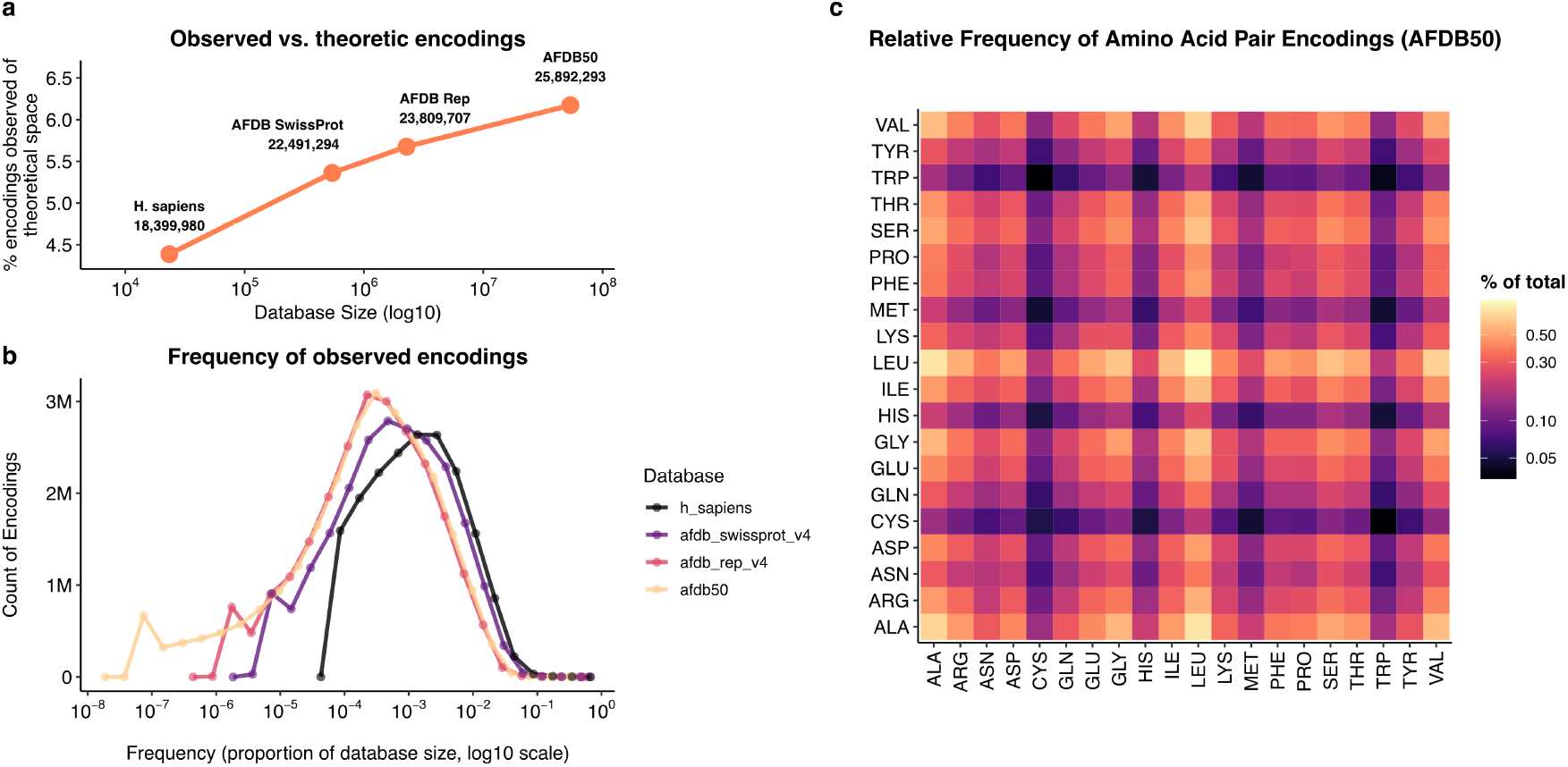
Sparsity and biological bias of Folddisco feature encodings. To characterize the usage of the feature space, we analyzed the distribution of encodings across four AlphaFold database subsets. Comparing the observed unique encodings to the theoretical 2^30^ limit (**top left**) reveals that *>* 93% of the space remains empty regardless of database size. This high sparsity justifies the implementation of a compressed index format. The frequency distribution of these observed encodings (**bottom left**) shows consistent patterns across databases, peaking in the 10^*−*4^–10^*−*3^ range. Additionally, the amino acid pair composition in AFDB50 (**right**) indicates that the index is dominated by hydrophobic, alpha-helical interactions (e.g., Ala-Leu), while Cysteine and Tryptophan pairs are under-represented.

**Supplementary Figure 2.**
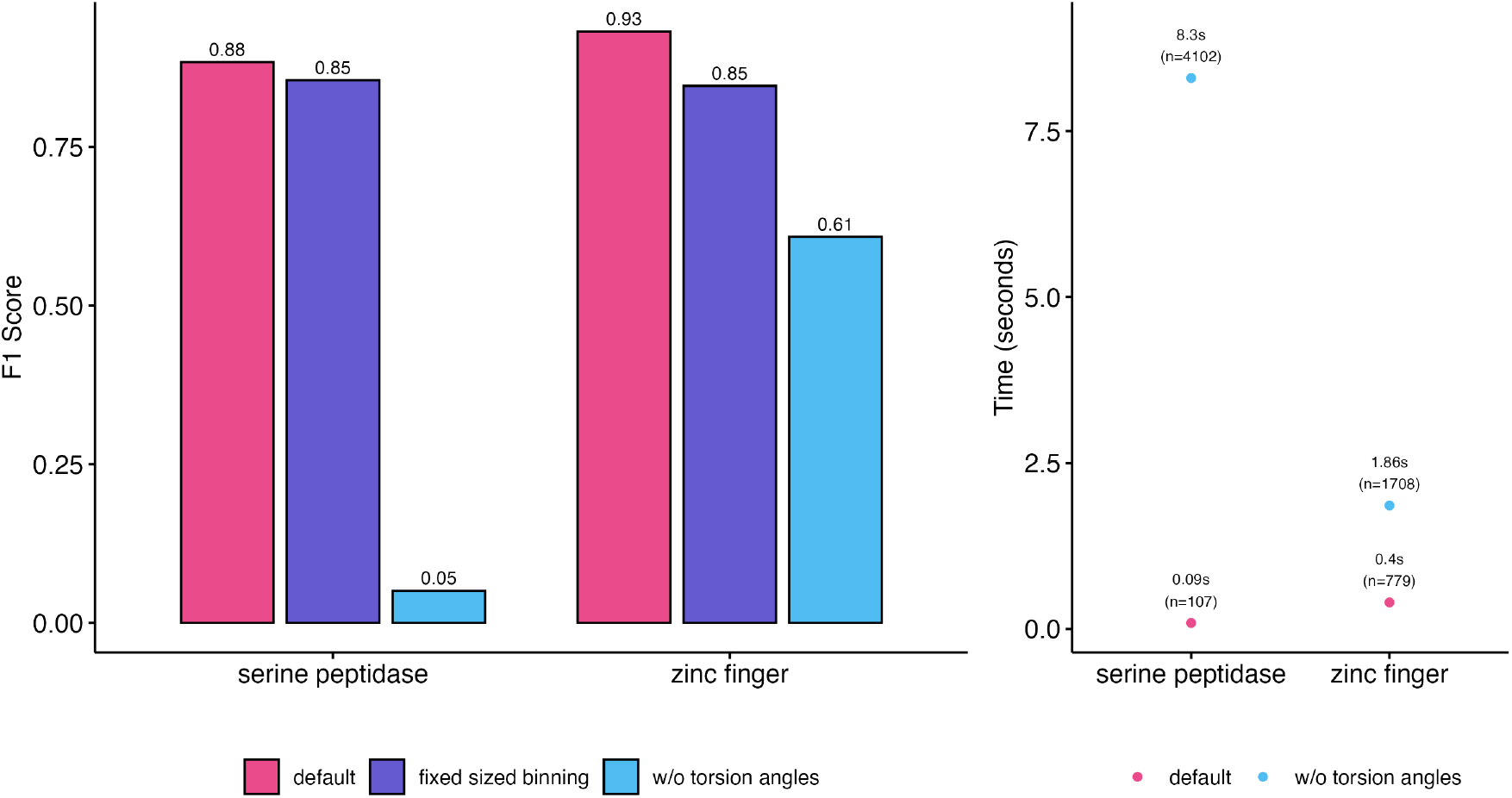
Effect of torsion angles inclusion and mode of encoding on Folddisco’s sensitivity. To evaluate the contribution of side-chain torsion angles, we performed similar analysis to that shown in **Fig. 1d**, querying the human fraction of the AFDB-proteome index for the serine peptidase and the zinc finger motifs. For each motif query, we conducted the analysis three times, using the tool’s default setting in which the torsion angles are included in the feature set and their cosine and sine values are discretized (pink), including them but discretizing the angles in radians directly (purple) or excluding them from the feature set (light blue). Excluding these angles results both in reduced sensitivity (F1 score, left panel) and increased runtime (right panel) due to an over-permissive prefilter step. Notably, the cosine and sine encoding strategy demonstrated an F1 gain in both queries compared to direct radian discretization.

**Supplementary Figure 3.**
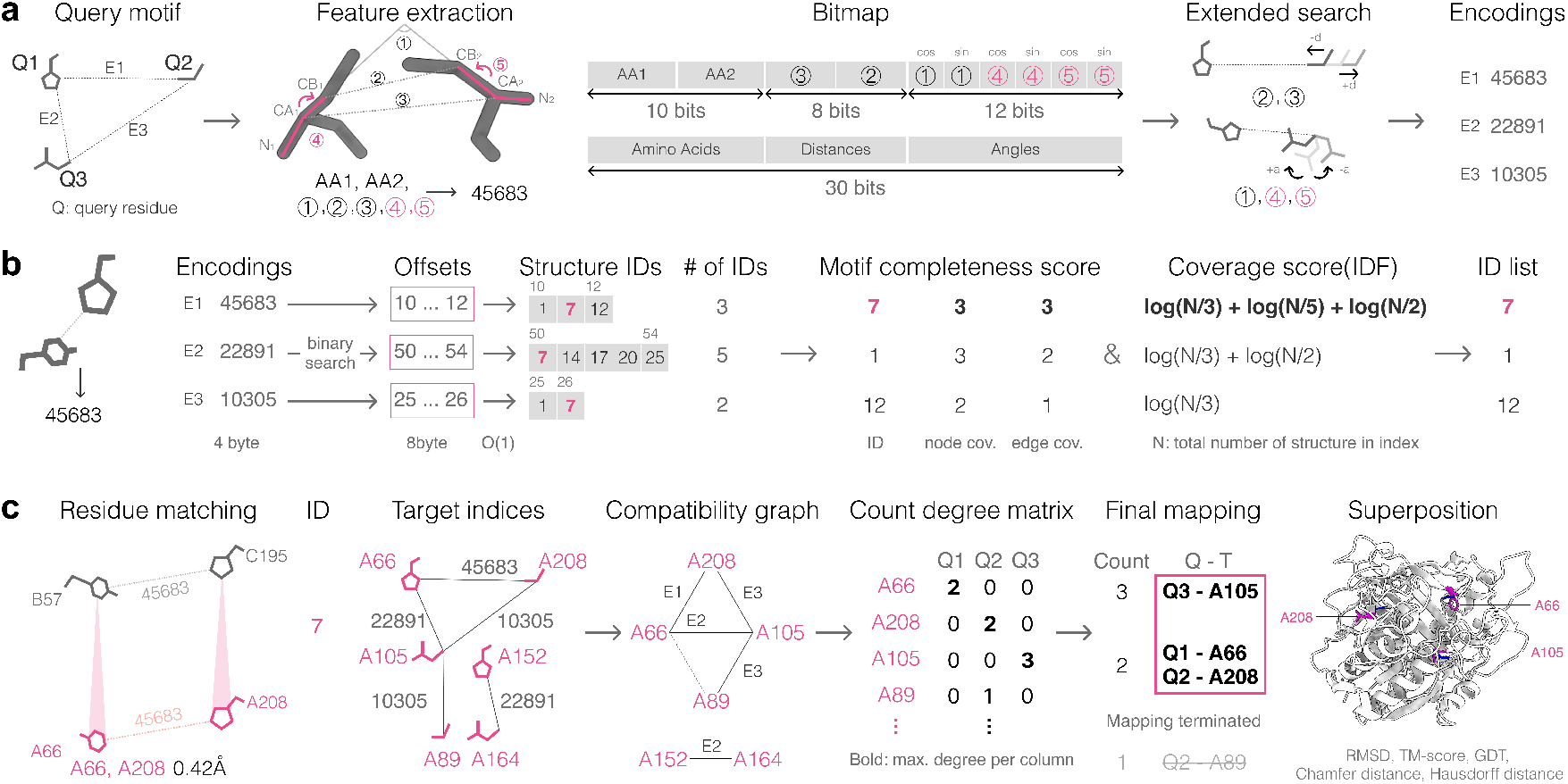
Folddisco’s workflow in detail. This figure extends **Fig. 1a-c** and illustrates the section “General workflow” of the Methods. **a**, Query feature extraction and feature encoding. From each pair of proximal (<20Å) query residues, Folddisco extracts a set of 7 features. Five of them (marked in black) were previously used by RCSB: amino acid identities (AA1, AA2), the angle between the Cα-Cβ vectors (circled 1), the distance between the Cβ atoms (circled 2) and the distance between the Cα atoms (circled 3); while two (marked in pink) are Folddisco-specific additions: the two dihedral angles in the atoms of N1-Cα1-Cβ1-Cβ2 (circled 4) and N2-Cα2-Cβ2-Cβ1 (circled 5). These features are encoded using a specific bitmap composition: 5 bits for amino acid identities, 4 bits for distances, and 4 bits for angles (allocated as 2 bits for cosine and 2 bits for sine values). Finally, to facilitate extended search, Folddisco generates up to 10 alternative feature sets by applying small deviations to individual features while keeping others fixed. **b**, Prefiltering Via binary search, Folddisco matches each encoding to a set of offset, which provides locations for structure IDs. For each structure ID, Folddisco computes scoring metrics that prioritizes structures with higher coverage (motif completeness score) and rare, functional motifs over ubiquitous secondary structures like helices or sheets (IDF-based coverage score). **c**, Residue matching For the structure ID with the highest coverage score, Folddisco matches target motifs into appropriate encodings. Folddisco constructs a compatibility graph, detects the connected component within the graph, and computes count degrees for each target. After computation, Folddisco implements final mapping correspondent to the count degree. Each query is mapped into the target in descending order - when all queries are mapped into a target, mapping is terminated. The mapping results would automatically be the proposed match to the motif that is superposed to it, and structure similarity metrics are computed.

**Supplementary Figure 4.**
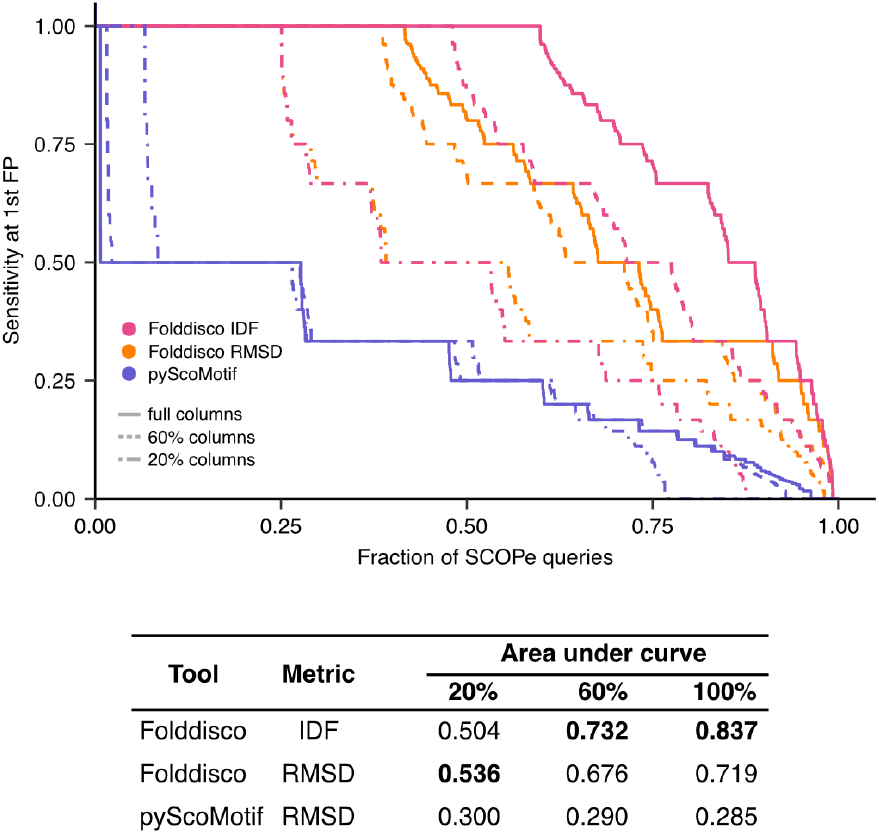
Sensitivity of Folddisco using two scoring modes and of pyScoMotif on SCOPe-constructed benchmarks. This figure extends **Fig. 1f**, where the goal is to match SCOPe sequences of the same family as the query before matching a different fold, using all conserved columns (“full”) or a random subsample of them (60%, 20%). Folddisco’s matches were ranked either by its IDF-based pre-filter coverage score (pink) or its full pipeline’s RMSD (orange), which is also used for ranking by pyScoMotif (blue). Folddisco achieves higher area under the curve values than pyScoMotif in all cases (bottom panel). Comparing Folddisco’s two scoring modes, RMSD is more sensitive for ranking matches of short queries (20% benchmark), while the IDF-based coverage score yields higher area the under curve values for the longer 60% and 100% benchmarks (bottom panel).

**Supplementary Figure 5.**
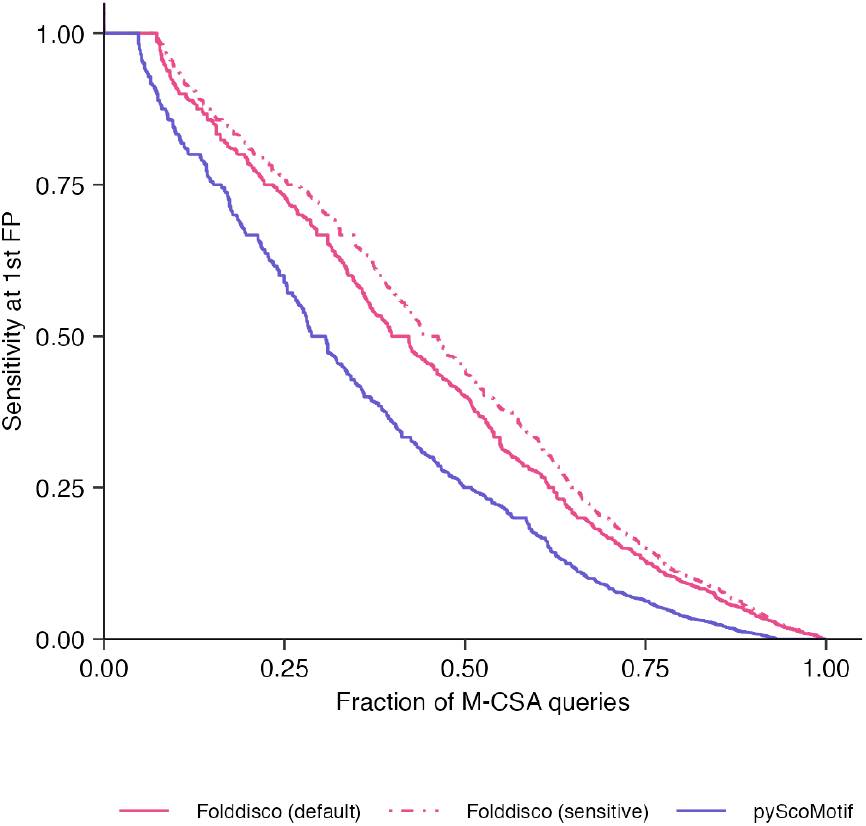
Sensitivity of Folddisco using two modes and of pyScoMotif on M-CSA-constructed benchmarks. This figure extends **Fig. 1g**. Here, we used a set of 713 queries from M-CSA reference catalytic sites, excluding motifs with only two residues to ensure comparability with pyScoMotif. The target database consisted of 62,122 PDB structures annotated as containing homologous catalytic sites. We measured sensitivity up to the first false positive (FP), defining a true positive as the retrieval of a PDB structure annotated as a homolog in M-CSA, with all remaining structures in the dataset serving as negatives. Folddisco (default), Folddisco (Sensitive) and pyScoMotif achieved AUCs of 0.432, 0.463 and 0.344, respectively.

**Supplementary Figure 6.**
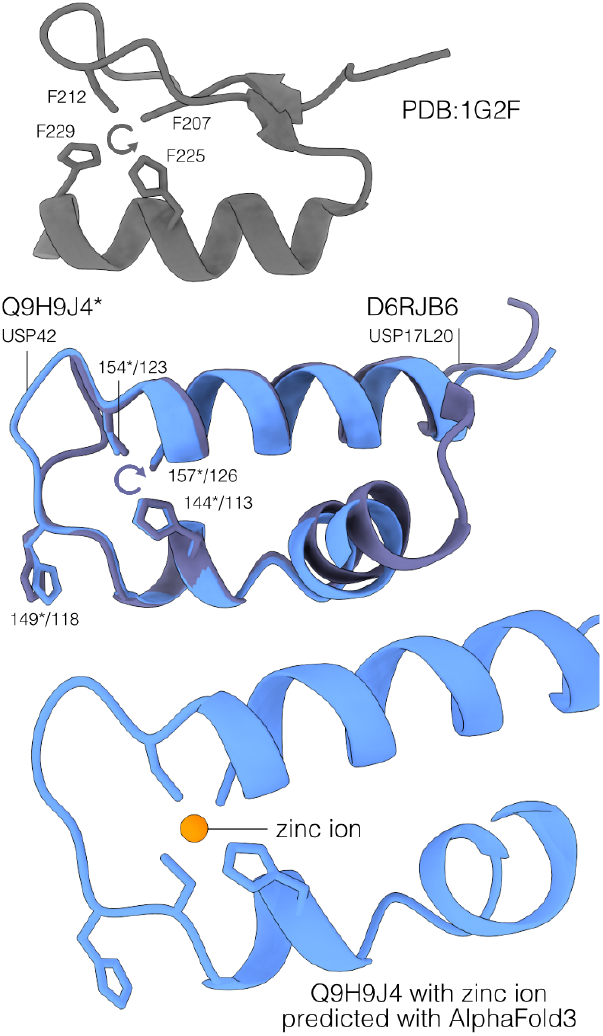
Partial zinc finger motifs identified in ubiquitin-specific peptidases. From *H. sapiens* proteome, Folddisco detected partial matches of the zinc finger motif (**top**) in ubiquitin-specific peptidases USP42 and USP17L20 (**middle**). Notably, the residue order is reversed in these matches. AlphaFold3 predictions confirmed zinc coordination within these motifs (**bottom**).

**Supplementary Figure 7.**
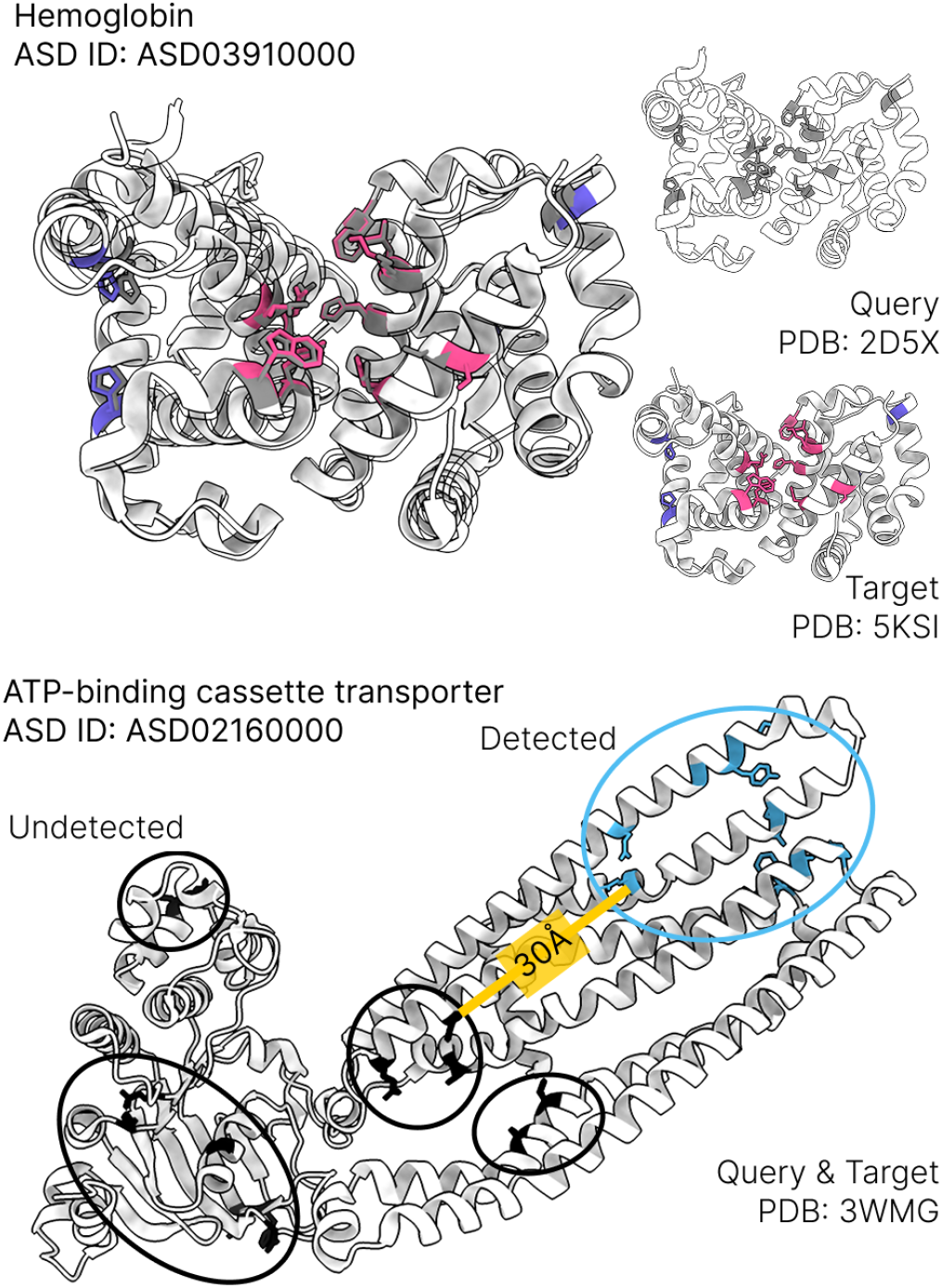
Evaluation of double-motif search capabilities. Proteins are often regulated by multiple distinct motifs, necessitating the simultaneous detection of functional sites. We assessed the capability to detect double motifs by querying 14 residues (10 allosteric, 4 active) in hemoglobin (**top**). Folddisco successfully identified matches sharing the same Allosteric Database (ASD) ID. Query residues are colored in gray, while matched active and allosteric residues are shown in purple and pink, respectively. In contrast, a test on an ATP-binding cassette transporter revealed a limitation in detecting widely distributed motifs (**bottom**). The algorithm failed to retrieve the distal allosteric sites (black) alongside the detected region (blue) because the spatial separation (>20 Å) exceeded the connectivity threshold.

**Supplementary Figure 8.**
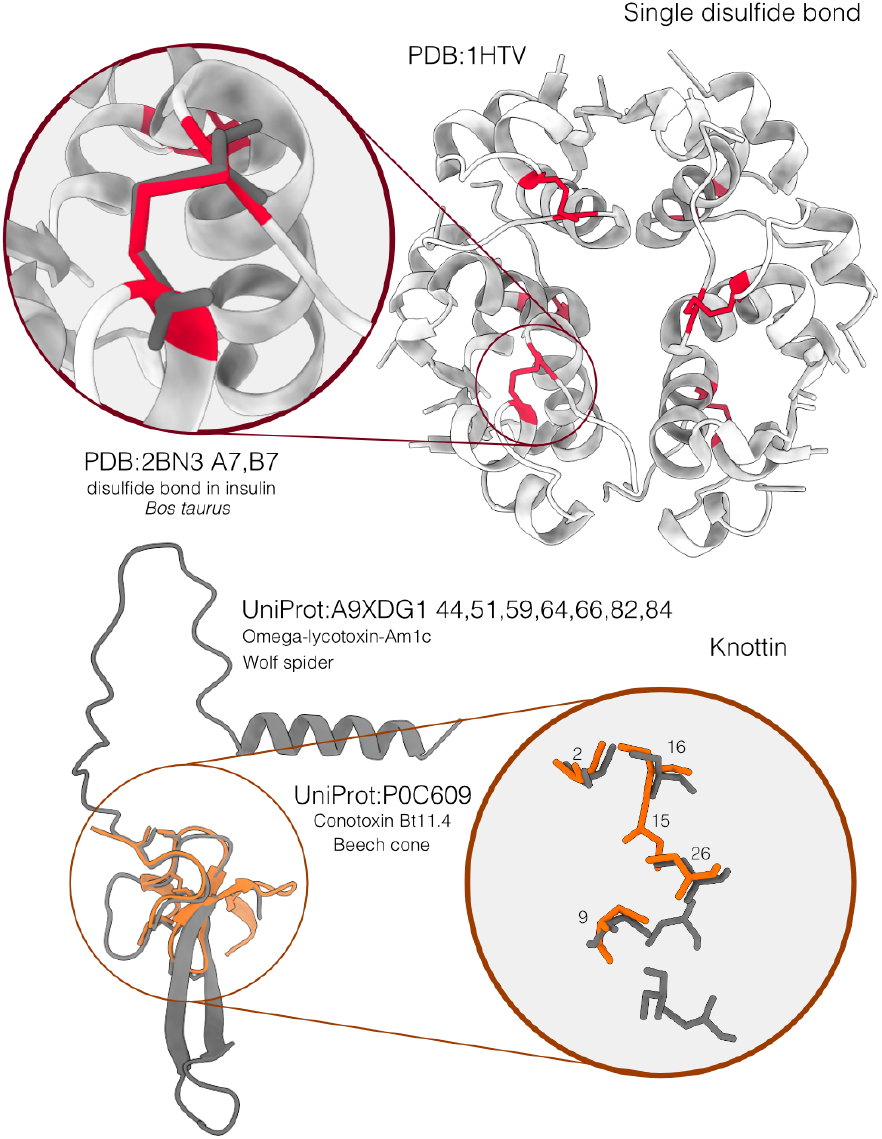
Detection of single disulfide bonds and knottin motifs. Folddisco identified single disulfide bonds (red) in insulin (**top**) and complex knottin motifs formed by multiple disulfide bonds (**bottom**). From search of a knottin motif in a spider toxin, a conotoxin was retrieved with a partial match.

**Supplementary Figure 9.**
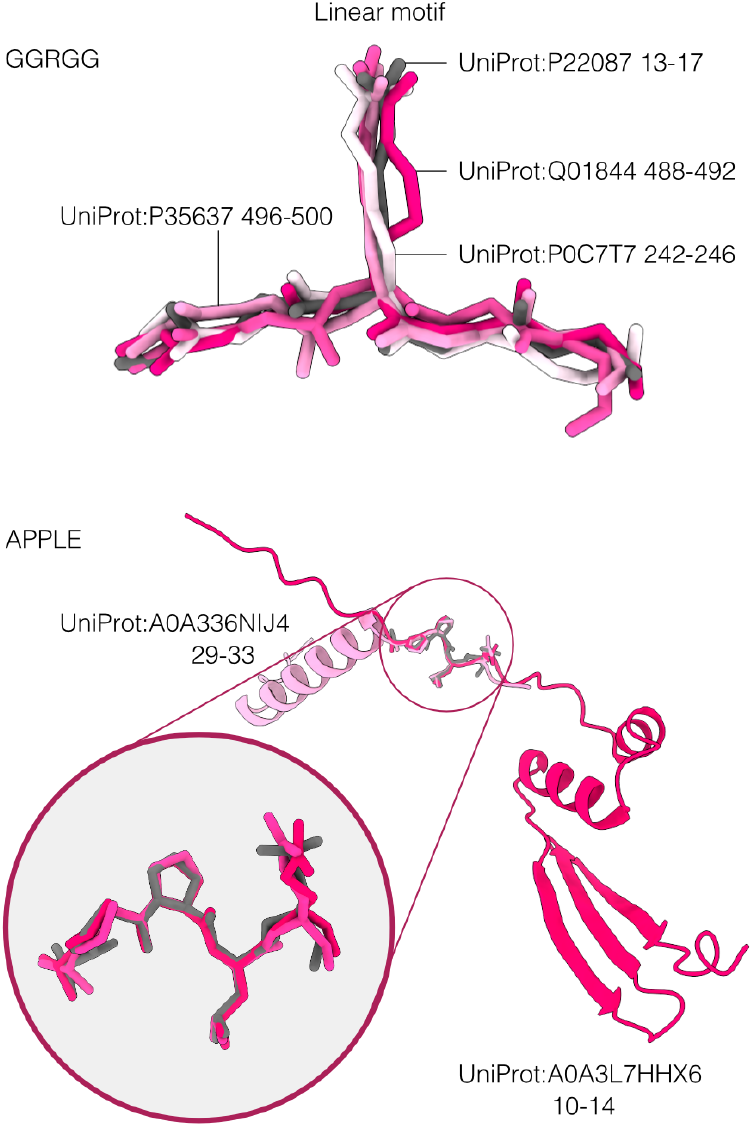
Identification of short linear motifs by Folddisco. Folddisco searched and detected the known “GGRGG” motif (**top**) and a arbitrarily generated “APPLE” motif (**bottom**). The query “APPLE” motif structure (gray) was predicted using ColabFold.

**Supplementary Figure 10.**
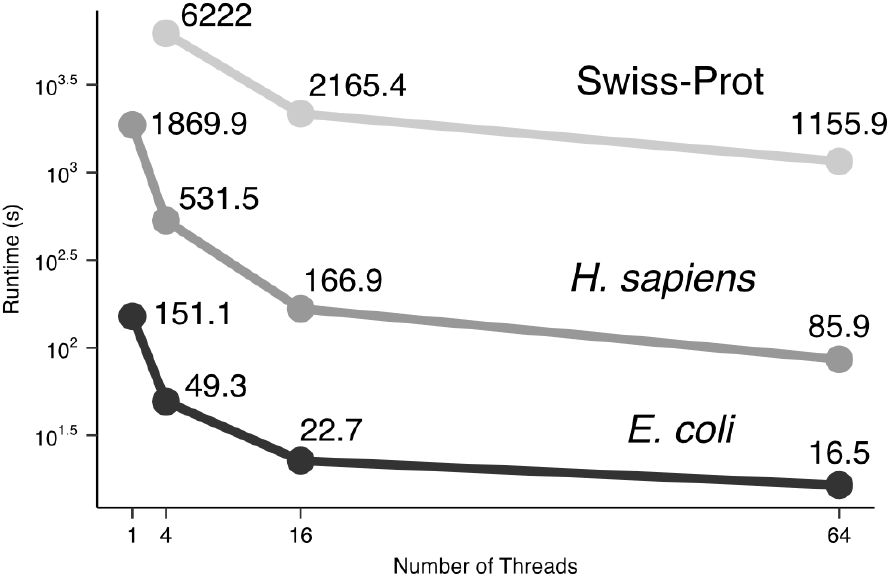
Runtime benchmarking of Folddisco index construction across databases and CPU cores. Index building time (in seconds) was measured for three databases (Swiss-Prot, human and *E. coli* subset of the AFDB-proteome) using 1, 4, 16, and 64 CPU cores.

**Supplementary Figure 11.**
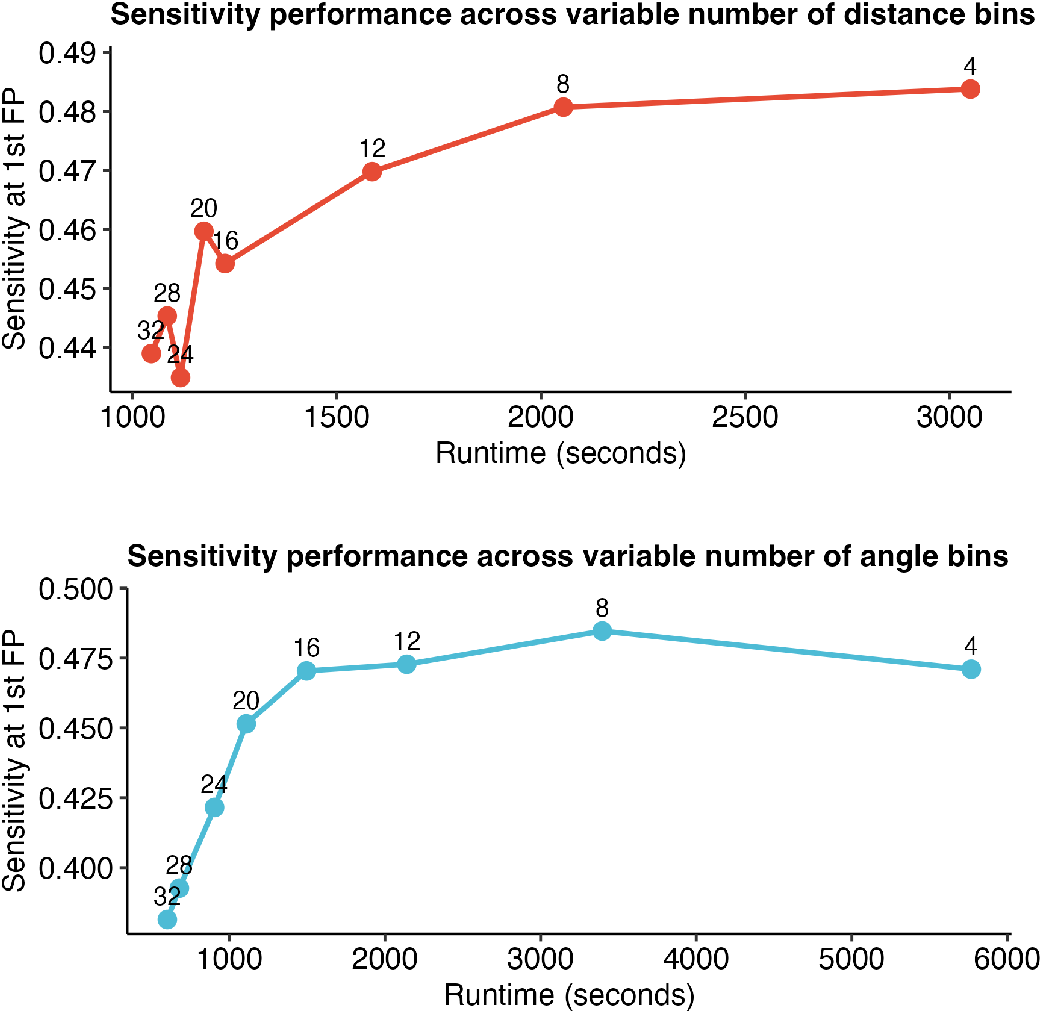
Sensitivity and runtime trade-off across number of bins. With M-CSA optimization subset, total runtime and sensitivity at the first false positive were measured for distance (**top**) and angle (**bottom**) discretization using 4 to 32 bins. While a higher number of bins generally improved speed at the cost of sensitivity, 16 bins yielded the optimal balance, maximizing sensitivity while maintaining a significant reduction in runtime.

**Supplementary Figure 12.**
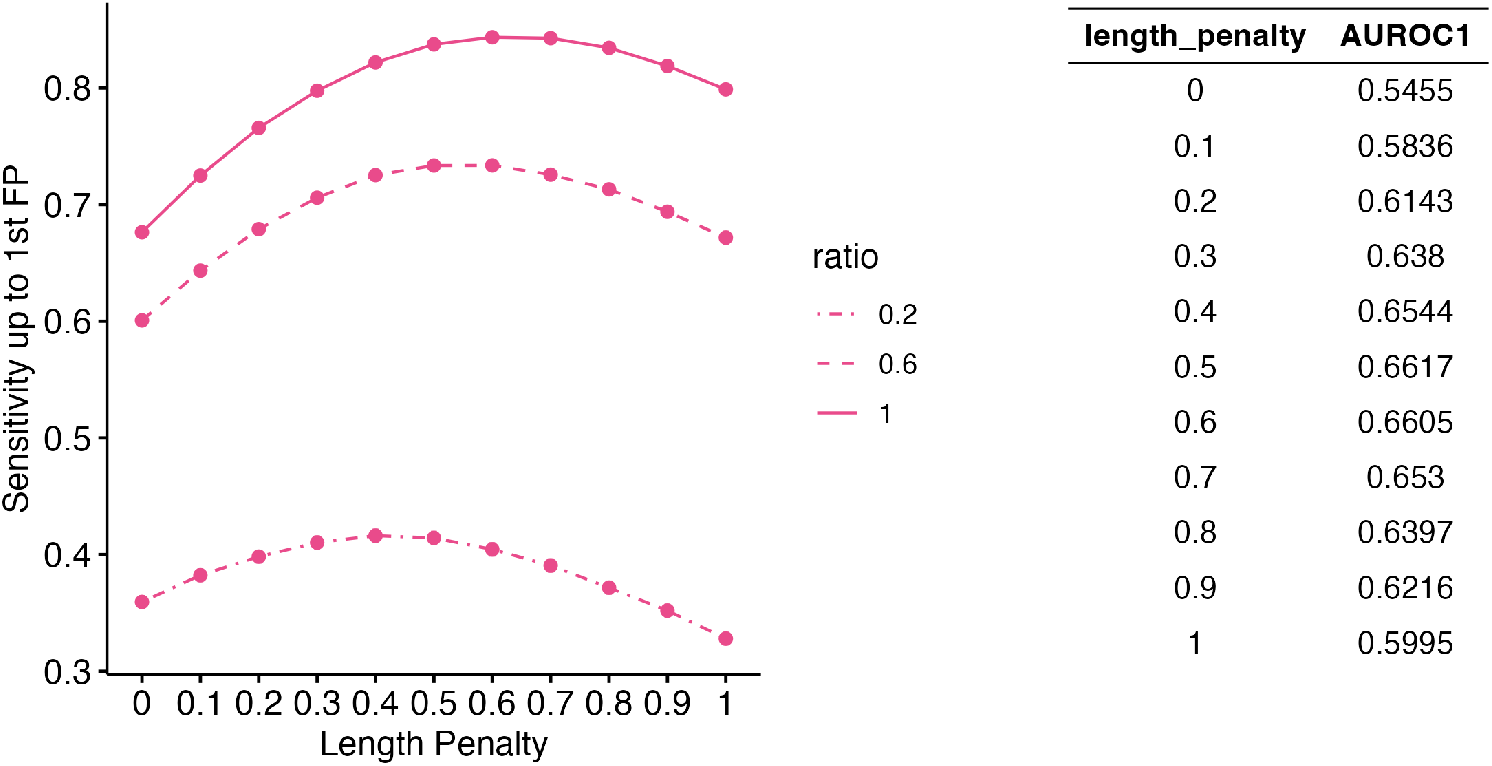
Tuning the length-penalty exponent. Using the SCOPe benchmark setting, sensitivity up to the first false positive of Folddisco pre-filter was evaluated as a function of the length penalty (0 to 1). **(left)** Sensitivity curves are shown for three SCOPe query sampling ratios. **(right)** The average sensitivity across these ratios. A length penalty of 0.5 yielded the highest average sensitivity and was identified as the optimal parameter.

